# Safety of Subdural Direct Current Stimulation: A Histological Study in the Ovine Brain Using Metal Electrodes

**DOI:** 10.64898/2026.07.13.737768

**Authors:** Michael Brosch, Hiroyuki Oya, Katherine N. Gibson-Corley, Oliver Flouty, Matthew A. Howard, Kirill V. Nourski

## Abstract

**Background:** Direct current (DC) stimulation can modulate neuronal activity in ways that differ from pulsatile stimulation, but its intracranial use has been limited by concerns about tissue injury at the electrode/tissue interface. Quantitative safety limits for DC delivered through metal electrodes directly to the brain remain poorly defined.

**Objective:** To estimate histological safety boundaries for DC stimulation delivered through metal electrodes in a large-brain gyrencephalic animal model.

**Methods:** Cathodal DC stimulation was applied to the exposed cortical surface of ten anesthetized sheep using platinum–iridium disc electrodes typically used in clinical applications (surface area ≤ 4.15 mm^2^). Currents of 5 to 1000 µA were delivered for 10 to 15 minutes at 36 cortical sites. Stimulation dose was quantified as charge density. Brains were removed shortly after stimulation and examined histologically for tissue damage, including necrosis, inflammation, gliosis, and demyelination. Lesion volumes were quantified and related to charge density.

**Results:** No lesions were observed at sites where no current or a low charge density (0.7 mC/mm^2^) was delivered. With stimulation, lesion probability and volume increased with charge density, although variability was substantial. Lesions occurred in 3 of 18 sites at lower charge densities (1.4 to 10 mC/mm^2^) and in 5 of 9 sites at higher charge densities (14.4 to 144.4 mC/mm^2^). Linear regression of lesion volume against charge density yielded an estimated zero-lesion intercept of 2.3 mC/mm^2^, whereas alternative nonlinear models predicted thresholds up to 8.7 mC/mm^2^.

**Conclusion:** These findings suggest that it may be possible to apply cathodal DC stimulation directly to the cortical surface through metal electrodes without detectable histological damage when current intensity, duration, and electrode size are appropriately constrained. These findings provide quantitative guidance for the safe application of DC directly to neural tissue in experimental and translational neuromodulation studies.

## Introduction

### Background

Direct electrical brain stimulation is a widely utilized invasive method for therapeutic, diagnostic and scientific purposes. It is successfully used in the form of current pulses, for example in cochlear implants to restore hearing [1] or in deep brain stimulation devices to manage conditions like Parkinson’s disease [2]. By contrast, interest in direct current (DC) stimulation has been limited, not only because pulsatile stimulation is well established, but also because of the challenges associated with safely delivering electrical charge for prolonged durations through metal electrodes into biological tissue. Nevertheless, the use of DC remains highly attractive as it affects nervous tissue in a distinct way [3, 4]. Pulsatile stimulation typically evokes action potentials via fast voltage-gated sodium channels in synchrony with the stimulation pulses. DC stimulation, in turn, alters the neuronal membrane potential, thereby modulating the efficacy of the synaptic inputs at different sites in generating action potentials. For this reason, DC can also suppress action potentials, which for current pulses must often be achieved indirectly, for example by activating inhibitory interneurons. Exploiting the specific physiological impact of DC could therefore offer new strategies in brain stimulation, particularly for challenges that are poorly addressed by pulsatile stimulation.

A key challenge in applying electrical currents directly to nervous tissue is avoiding potentially harmful side effects at the electrode–tissue interface. Violating the safe charge injection threshold can result in pH changes, electrode corrosion, toxic byproducts, gas formation, and heat production [5-7]. Toxicity issues can be partially circumvented by using salt bridges to buffer the electrode-tissue interface, although such bridges are often impractical for intracranial applications [6]. For metal electrodes, damage can be reduced by applying brief charge-balanced pulses on the order of microseconds to milliseconds per phase. For platinum electrodes, a commonly cited safety threshold below which no tissue damage occurs during prolonged pulsed stimulation is 0.3 µC/mm^2^ per stimulation phase for electrode surfaces on the order of several square millimeters [5, 8–10].

Charge balancing is not feasible with DC stimulation, making it substantially more challenging to mitigate potentially harmful electrochemical processes [6, 11, 12]. However, a few studies (see below) suggest that very low DC doses may not cause adverse effects in the neural tissue. Despite this, the threshold values for safe DC application remain poorly defined.

### Previous studies on DC safety

In a seminal *transcranial* DC stimulation (tDCS) study on rats, Liebetanz et al. [13] applied monopolar cathodal DC for 3 to 270 minutes directly to the skull using an electrode with a surface area of 3.5 mm^2^. Their results established that *charge density*-the product of amplitude and duration of current application divided by electrode surface area - is a meaningful parameter to describe the relationship between DC dose and the extent of tissue damage and for safety assessment (see also [14,15]). The study further revealed a clear linear relationship between the two parameters, with a strong coefficient of determination (*R*^2^ = 0.94) which allowed extrapolation of a zero lesion size intercept of 52.4 mC/mm^2^ below which tDCS should be safe. Importantly, subthreshold effects of DC did not accumulate over consecutive days, possibly because detrimental effects of tDCS accumulate sufficiently slowly to be cleared by diffusion and the brain’s interstitial fluid homeostasis mechanisms [16] before reaching toxic concentrations. The existence of a histological safety limit, possibly at an even higher value, was later confirmed by Zhang et al. [17]. A similar safety limit was also reported for anodal stimulation [4, 15].

Histological safety limits also seem to exist for DC directly delivered to nervous tissue. This was observed for chronic DC stimulation applied through metal electrodes with a size of 2 mm^2^ implanted in the thoracic spinal cord of rats [18]. Histological analyses revealed that delivery of 1.5 μA for ten days resulted in no tissue changes in any of the five animals studied under this condition, whereas such changes were detectable in many of the remaining 30 animals at current levels between 3 and 50 μA, including cavitations, inflammatory responses, fibrosis, and gliosis. Further support for a dose-dependent histological boundary comes from observations that lesion size decreased with decreasing stimulation dosage in this and other studies [19-22], the latter employing larger currents (up to 2.4 mA), shorter durations (seconds to minutes), similar and smaller electrodes (including microelectrodes), and targeting other nervous structures (cortex, cerebellum and sciatic nerve) in various species (pig, sheep, monkeys and mice). A linear extrapolation of the decreasing trends reported by Bray and colleagues [22], for example, indicates that no tissue damage would occur below a total charge of 2.9 mC and no cavitations would occur below 10.4 mC.

An alternative way to estimate safety for invasive DC stimulation is to assess functional properties of nervous tissue. Such studies have revealed stimulation conditions under which no measurable functional impairments were detected. Bray et al. [22] continued to record reliably similar spike waveforms from the cortex of monkeys even if DC was repetitively applied at levels that produced histological damage. A charge density above 16 mC/mm^2^ was reported for 15-minute delivery of DC (± 1mA) to the cortical surface of monkeys [23] without impairments in spiking activity or local field potentials in the underlying cortex. Functional no-effect levels have also been reported for DC delivered through metal electrodes to the semicircular canal in the inner ear [12] and to the sciatic nerve [11]. Finally, Hurlbert and colleagues [18] reported impairments of motor functions controlled by the spinal cord at current levels above 18 μA, which were substantially higher than those at which histological changes were seen (≥3 μA). Together, these findings suggest that histological alterations may occur at stimulation levels below those required to produce detectable deficits in the functional measures assessed.

### Aim of the present study

Here, we aimed to estimate safety boundaries for DC applied directly to nervous tissue through metal electrodes. Current knowledge is limited to a very small set of stimulation parameters, electrode sizes and specific neural structures. To this end, we applied DC directly to the cortical surface of sheep, a large animal gyrencephalic model, and examined the cortical tissue beneath the stimulation site for histopathological abnormalities, including demyelination, inflammation and necrosis. We used DC parameters and electrodes with potential relevance for neurocognitive and other research purposes in animals and humans [24].

## Material and methods

Ten adult sheep were used in this investigation which was reviewed and approved by the University of Iowa Animal Care and Use Committee (IACUC# 0902039). Surgical details are described in Gibson-Corley et al. [25] and Flouty et al. [26]. The animals were initially anesthetized with Isoflurane inhalation and then maintained throughout the surgical procedure through an endotracheal tube. Isoflurane was discontinued approximately 30 minutes prior to the intracranial direct current stimulation (iDCS) experiments. The anesthesia was maintained throughout the DC stimulation with intravenous injection of Propofol (0.4 mg/kg/h), and virtually zero % Isoflurane concentration was confirmed before the stimulation. Blood pressure, oxygen saturation (SpO_2_), and body temperature were monitored continuously during the surgical and experimental procedure. A right-sided craniotomy (approximately 10 × 10 cm, bilateral in one sheep) was performed for each animal using a high-speed drill. The dura was incised and reflected, such that the brain surface was exposed while leaving the pia intact. A total of 30 ml of lidocaine (2%) was injected at the incision sites for the craniotomy and durotomy. Corneal reflex was tested periodically to ensure proper depth of anesthesia, and body temperature was maintained within normal limits using a heating pad and warm intravenous saline infusions, as needed.

Four-contact strip electrodes (platinum-iridium, disc shaped, exposed diameter = 2.3 mm, and 1 mm in animal S1; corresponding surface areas: 4.15 mm^2^ and 0.79 mm^2^, center-to-center distance between electrodes: 1 cm; Ad-Tech Medical Instrument Corporation, Oak Creek, WI, USA) were used for current delivery (**Figure 1**). Negative (cathodal) iDCS was applied through individual electrodes directly placed on the brain surface from where the current returned to a subgaleal strip at the vertex. Proper contact with the brain surface was ensured by placing wet neurosurgical pads on the back side of the stimulating electrodes. A TDT IZ2-H stimulator (Tucker-Davis Technologies, Alachua, FL, USA) was used for current delivery. iDCS was applied for 10 minutes (15 minutes in animal S2). Current intensities, stimulation duration and lesion size are summarized in **Table 1** for 36 sites across ten sheep. These stimulation parameters were chosen because comparable parameter ranges may be relevant for neurocognitive studies in animals and for translational studies in human epilepsy patients temporarily implanted with similar electrode arrays [24]. Negative currents have predominantly suppressive effects, enabling causal assessment of functional roles of spatially confined brain regions. Stimulation duration would allow repetitive testing of iDCS effects across many (~100) behavioral task epochs, lasting several seconds each.

**Table 1.** Summary of all stimulation experiments (36 cortical sites in 10 sheep).

| Sheep ID | Experiment # | Current (μA) | Charge density (mC/mm<sup>2</sup>) | Lesion volume (mm<sup>3</sup>) |
| --- | --- | --- | --- | --- |
| S1 | 1 | 50 | 38.2 | 0 |
|  | 2 | 70 | 53.5 | 0 |
| S2 | 1 | 300 | 65.0 | 0 |
| S4 | 1 | 150 | 21.7 | 1.76 |
|  | 2 | 50 | 7.2 | 0 |
|  | 3 | 300 | 43.4 | 0.81 |
| S5 | 1 | 10 | 1.4 | 0 |
|  | 2 | 25 | 3.6 | 0 |
|  | 3 | 50 | 7.2 | 0 |
| S6 | 1 | 25 | 3.6 | 0 |
|  | 2 | 50 | 7.2 | 0 |
|  | 3 | 100 | 14.4 | 0 |
| S7 | 1 | 50 | 7.2 | 0 |
|  | 2 | 25 | 3.6 | 0 |
|  | 3 | 0 | 0 | 0 |
|  | 4 | 100 | 14.4 | 0.42 |
| S9 | 1 | 50 | 7.2 | 1.57 |
|  | 2 | 0 | 0 | 0 |
|  | 3 | 10 | 1.4 | 0.09 |
|  | 4 | 25 | 3.6 | 2.12 |
| S10 | 1 | 50 | 7.2 | 0 |
|  | 2 | 50 | 7.2 | 0 |
|  | 3 | 5 | 0.7 | 0 |
|  | 4 | 10 | 1.4 | 0 |
| S14 | 1 | 1000 | 144 | 5.50 |
|  | 2 | 0 | 0 | 0 |
|  | 3 | 0 | 0 | 0 |
|  | 4 | 0 | 0 | 0 |
|  | 5 | 1000 | 144 | 4.71 |
|  | 6 | 0 | 0 | 0 |
|  | 7 | 0 | 0 | 0 |
|  | 8 | 0 | 0 | 0 |
| S15 | 1 | 25 | 3.6 | 0 |
|  | 2 | 25 | 3.6 | 0 |
|  | 3 | 25 | 3.6 | 0 |
|  | 4 | 25 | 3.6 | 0 |

**Figure 1.**
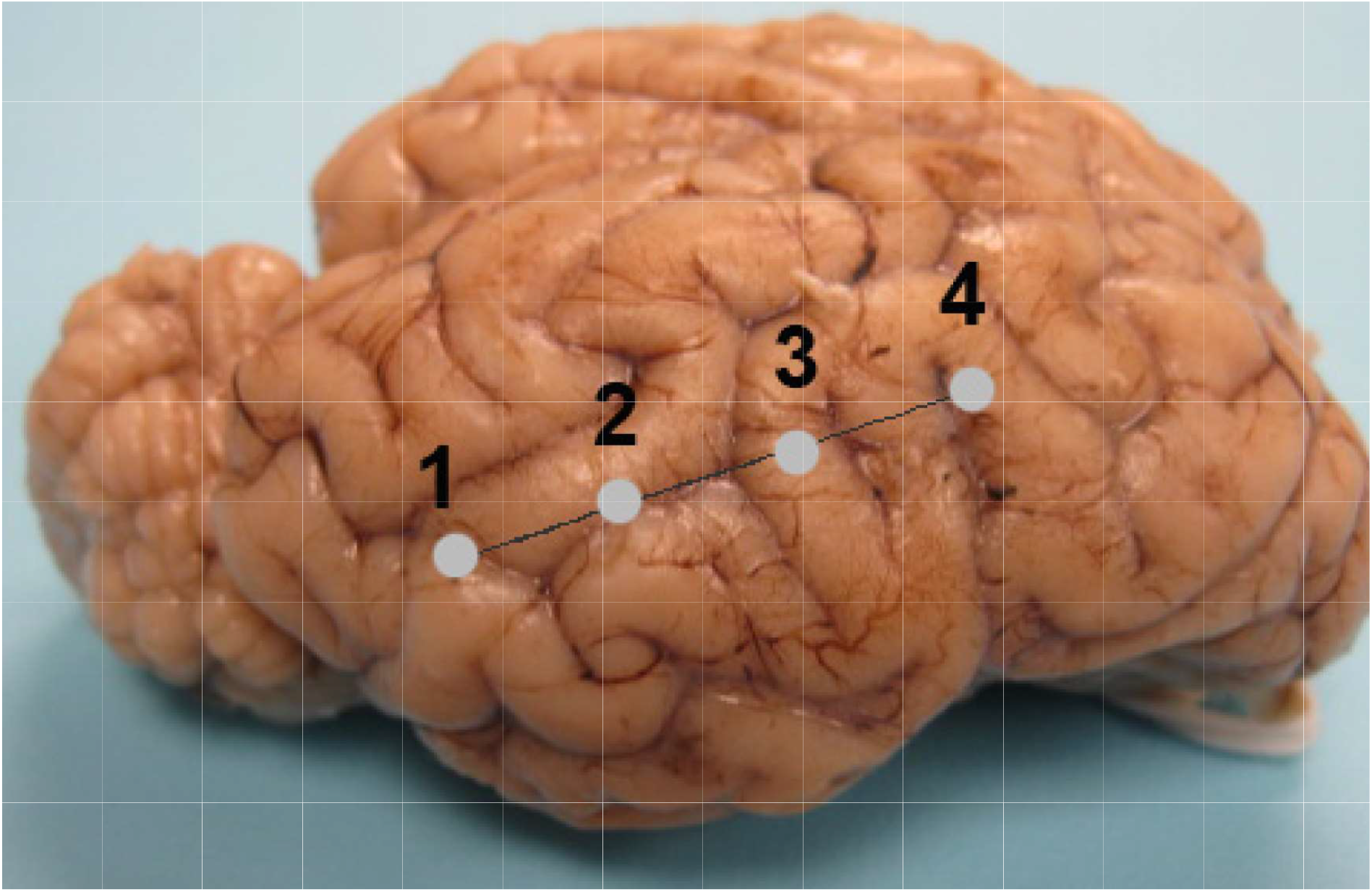
Photograph of the right hemisphere of a sheep brain (S7) showing the location of four disc electrodes. Each electrode was individually used as cathode for monopolar intracranial direct current stimulation. Distance between electrodes was 1 cm.

All stimulation sites were located in the right hemisphere, except for sites 5-8 in sheep S14, which were in the left hemisphere. Total charge density *J* was the product of intensity *I* and duration t of current delivery, divided by the electrode surface size A: *J* = *I*t/A*.

The experiments lasted approximately 8 hours. Following the approved protocol, animals were euthanized by an injection of a lethal dose of KCl while still under propofol anesthesia approximately 5 minutes after the experiment. After that, craniotomies were performed to gently remove the brain from the skull (both cerebral hemispheres) for fixation. Whole brains were immersed in 10% neutral buffered formalin for fixation at room temperature on constant agitation for approximately 5 days, followed by trimming of the brain with a focus on areas that underwent iDCS. These samples were cassetted and then secondarily fixed with 10% neutral buffered formalin for another 48-72 hours. Samples were processed using an automated tissue processor (Thermo Excelsior ES Tissue Processor, Waltham, MA, USA) on a long processing program for sufficient dehydration, embedded, sectioned at 4 µm, and stained with hematoxylin and eosin to identify neural inflammation and Luxol fast blue to identify demyelination for histological examination by a board-certified veterinary pathologist (KGC). No lesions were seen below the subgaleal strip at the vertex.

To calculate the electric field generated by monopolar stimulation through an electrode placed on the pia, we modelled the electrode as a circular area placed on a semi-infinite, homogeneous conductor [27]. Outside the electrode, the current flow can be approximated as point-source / half-space-like, and the electric field strength E decreases orthogonally to the brain surface with distance z according to

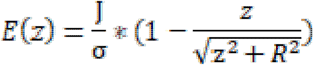

For an electrode radius R of 1.125 mm, a tissue conductivity *σ* of 0.2 mS/mm (average σ of gray and white matter; [28]) and a stimulation current *I* of 15.9 μA (which after 10 minutes accumulates to a charge of 9.54 mC and a charge density *J* of 2.3 mC/mm^2^), the resulting electric field strengths are 20.00 mV/mm at *z* = 0, 6.71 mV/mm at *z* = 1 mm, 2.57 mV/mm at *z* = 2 mm, and 0.49 mV/mm at *z* = 5 mm.

Parallel to the cortical surface, the electric field strength decreases with distance r from the electrode according to

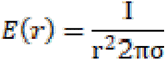

At the edge of the electrode, the electric field strength is 20.0 mV/mm. It decreases to 8.79 mV/mm at r = 1 mm from the edge, and to 4.63 mV/mm at r = 2 mm from the edge. Changes in current strength lead to proportional changes in electric field strength.

## Results

An example of a lesion following 100 µA iDCS in sheep S7 is shown in **Figure 2**. A focus approximately 1.8 mm in length, 0.5 mm deep and 0.6 mm wide (corresponding to a lesion size of 0.42 mm^3^, calculated using the formula for the volume of a cylinder with given length, depth and width) is characterized by a superficial zone of hypereosinophilia with loss of normal cellular and nuclear structures interpreted as acute liquefactive necrosis (zone 1). Deep to this zone is a band of tissue with numerous deeply basophilic, pyknotic nuclei (coagulation necrosis) with loosening of the neuropil (zone 2) (**Figure 2A, B**). There is increased white space surrounding blood vessels in the deeper tissues. Diffusely throughout the leptomeninges there is a moderate to marked increase in neutrophils, both free within the meninges and marginating within meningeal blood vessels. Meningeal blood vessels are also mildly congested (**Figure 2A**). At the junction of gray and white matter, multiple deeply eosinophilic to basophilic, angular neurons (necrotic neurons) are present, often bounded by multiple neuroglial cells (satellitosis). There also appears to be a moderate increase in neuroglial cells in this zone (gliosis) (**Figure 2C**).

**Figure 2.**
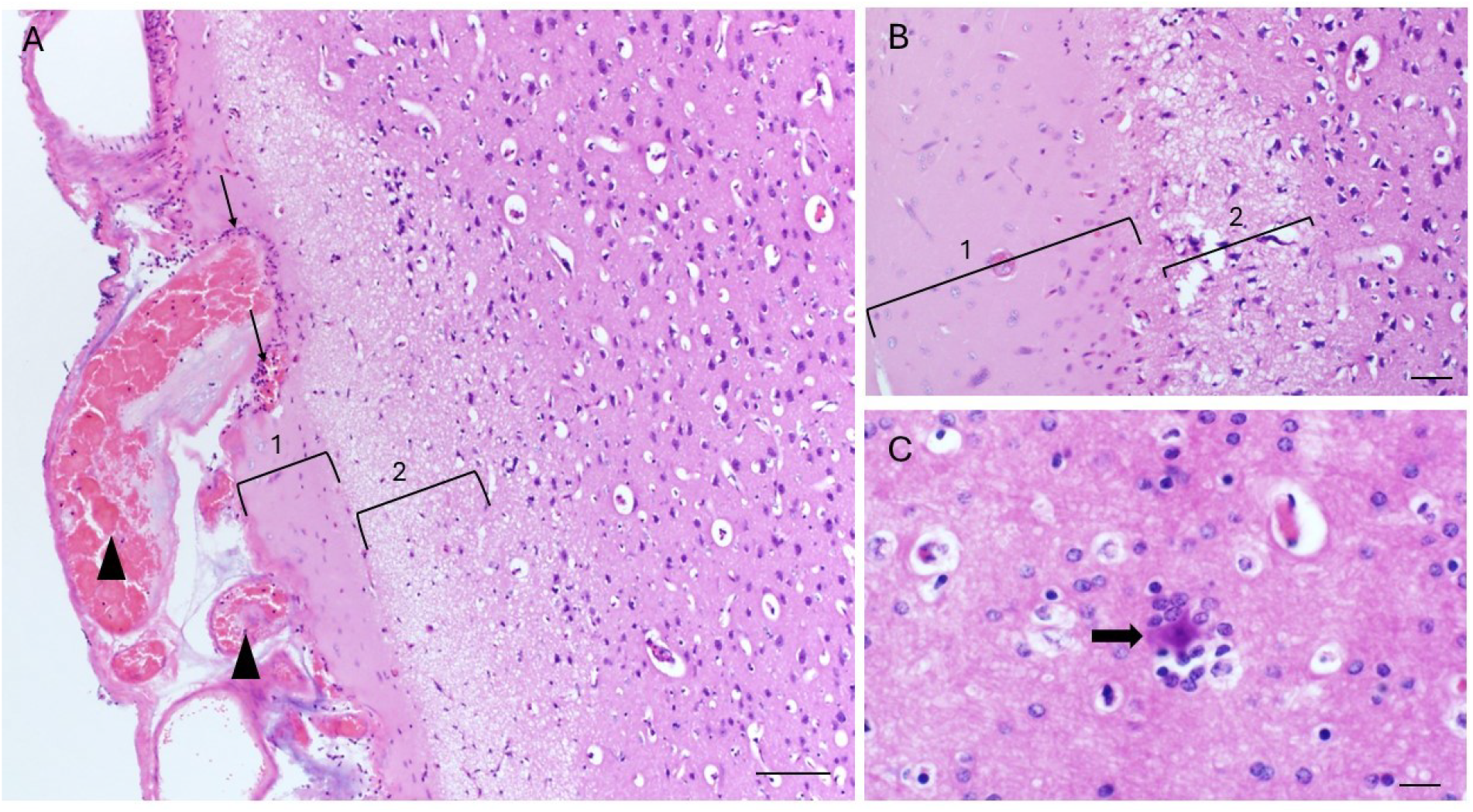
HE-stained (Hematoxylin and Eosin) photomicrographs of brain sections from sheep S7. **A)** Low magnification view of a focal lesion characterized by a superficial zone of hypereosinophilia with nuclear pyknosis (zone 1) with a deeper zone of rarefaction or loosening of the neuropil characterized by increased clear space (zone 2). Also noted was congestion of meningeal vessels (arrowhead) and perivascular neutrophils (thin arrows). Bar = 100 µm. **B)** Higher magnification of the lesion defining zones 1 and 2. Bar = 50 µm. **C)** High magnification image deep to zones 1 and 2 showing neuronal necrosis (thick arrow) with satellitosis. Bar = 20 µm.

The relationship between charge density and lesion size is shown in **Figure 3** for all 36 cases from 10 sheep. When no current was delivered, no lesions were observed in any of the 8 cases, indicating that neither electrode placement nor iatrogenic manipulations during surgery and craniotomies caused detectable tissue damage. In contrast, when current was delivered, both the probability of observing a lesion and its size increased with charge density, despite considerable variability across cases tested with the same and different charge densities.

**Figure 3.**
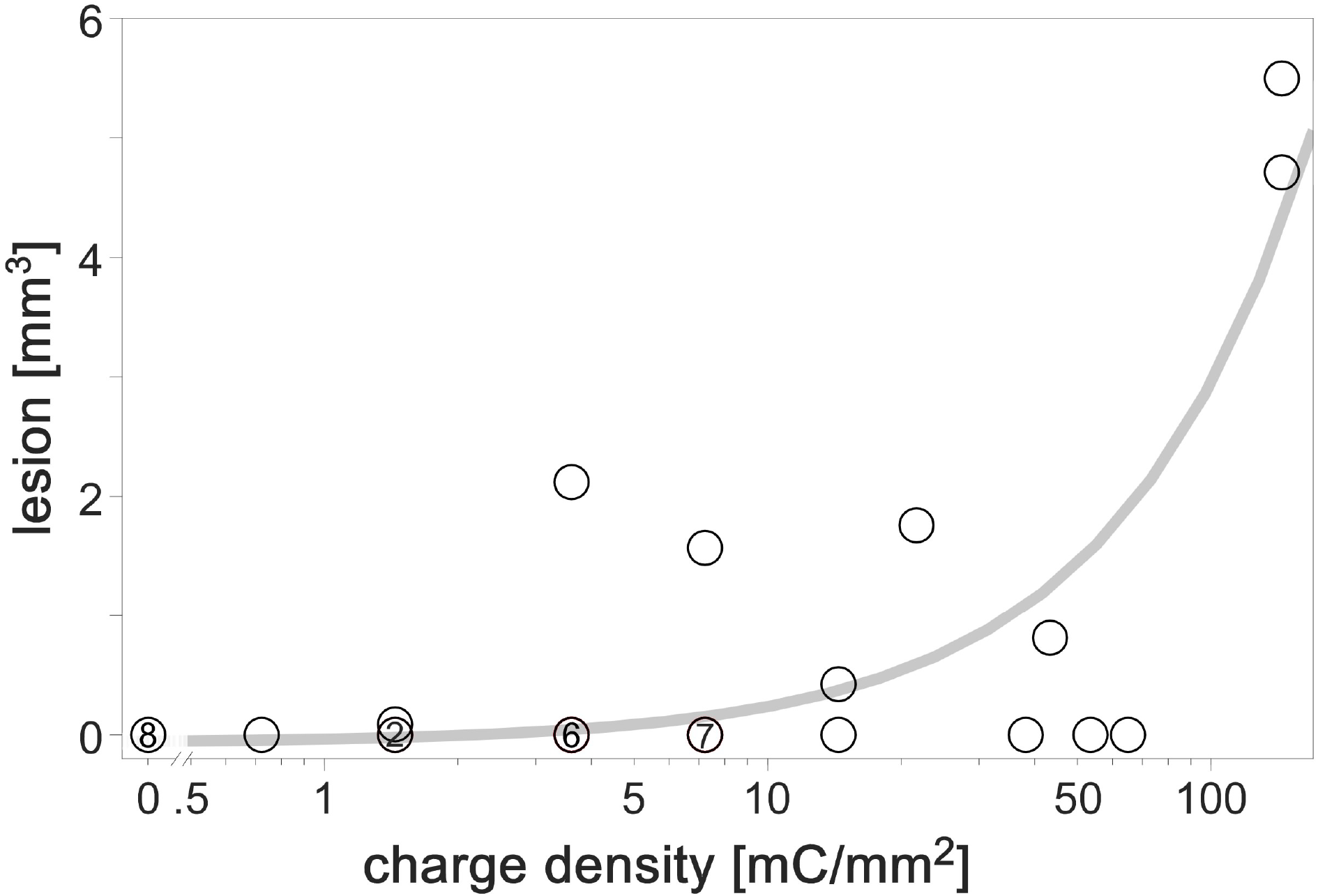
Relationship between charge density and lesion volume. A semilogarithmic representation was used to facilitate visualization across the full range of charge densities and lesion volumes. Numbers within circles indicate cases where multiple observations were made. The gray line shows the linear regression in this semilogarithmic plot. It has a coefficient of determination *R*^2^ of 0.68 and intersects the abscissa at a charge density of 2.3 mC/mm^2^, suggesting that no lesion is generated below this value.

At the lowest charge density (0.7 mC/mm^2^), no lesion was observed. At moderate charge densities (1.4 to 10 mC/mm^2^), lesions were observed in 3 out of 18 cases (17 %). This percentage increased to 56 % (5 out of 9 cases) at higher charge densities (14.4 to 144.4 mC/mm^2^). To quantitatively estimate the relationship between charge density and lesion size, a linear regression analysis was performed, following the approach of Liebetanz et al. [13] and yielding a coefficient of determination (*R*^*2*^) of 0.68. The regression line intersected the abscissa at a charge density of 2.3 mC/mm^2^, predicting that no lesion is generated below this threshold. Alternative model fits were also explored. Concave relationships between lesion volume and charge density, (e. g., with power-law models with exponents < 1) yielded lower *R*^*2*^ values and predicted smaller threshold charge densities. In contrast, convex relationships (e. g., exponents > 1) could produce higher *R*^*2*^ values, reaching a maximum of 0.77 for an exponent of 2.28, and predicted a substantially higher threshold charge density of 8.7 mC/mm^2^. In conclusion, depending on model assumptions, our data suggest a conservative zero-lesion value near 2.3 mC/mm^2^, with alternative nonlinear fits indicating that higher thresholds may also be plausible.

## Discussion

To our knowledge, this is the first controlled study in a large-animal model addressing safety issues associated with delivering direct current to the cortical surface. We used sheep because their brain size and gyrencephalic organization share relevant anatomical features with the human brain. Our findings indicate that, within the resolution of the present histological methods, there is a dose below which tissue damage was not detected. This suggests that intracranial DC delivery is feasible when current intensity, duration, and electrode size are appropriately constrained. Our study extends previous observations that, under specific conditions, DC can be delivered to neural tissue without causing histological damage [18] or detectable impairment of neural function [11,12, 22, 23]. Therefore, the specific effects of DC stimulation may not only be exploited transcutaneously and transcranially, but also intracranially.

In the present study, suprathreshold cathodal DC delivered for 10–15 minutes to the pial surface of the ovine brain resulted, within minutes after stimulation, in localized coagulative necrosis, increased neutrophil presence, necrotic neurons, satellitosis, and gliosis in the underlying brain tissue. These findings are broadly consistent with previous reports of the effects of suprathreshold DC stimulation, which include coagulative necrosis, parenchymal rarefaction, perivascular microhemorrhage, narrow leukocytic transition zones, myelin and axonal degeneration, fibrosis, and gliosis [13, 15, 17, 18, 20-22]. In contrast to some prior reports, however, we found no evidence of cavitation.

Differences in the severity and nature of tissue damage of suprathreshold DC across studies likely arise from multiple factors. These include current direction, intensity, and duration, as well as properties of the electrode-tissue interface, including electrode material, geometry, coupling medium, and placement. While no differences in thresholds were found between cathodal and anodal tDCS [13, 15, 17], Horsley and Clarke [19] reported different histological damage following anodal and cathodal stimulation. Hurlbert et al. [18] found more severe tissue damage near the anode than the cathode, suggesting that the safety values identified in the present study may be lower for anodal DC stimulation. Additional factors include post-stimulation survival time (minutes in the present study, hours in Bray et al. [22], and days in Hughes et al. [20] and Hurlbert et al. [18]), the relative distribution of gray and white matter, cell-type composition and spatial organization, and the histological methods used to assess tissue damage (e.g., hematoxylin-eosin, Luxol fast blue, and glial fibrillary acidic protein staining). Another important consideration is the extent to which DC-induced neural damage can be distinguished from damage caused by mechanical manipulations during surgery, electrode placement, and the foreign body response. The absence of detectable histological alterations at lower doses does not necessarily indicate that electrochemical reactions were absent. Rather, it indicates that, under the tested conditions, such reactions did not produce tissue damage detectable with the histological approach used here.

Several mechanisms have been proposed to explain how electrical stimulation can cause brain tissue damage [5]. One class of mechanisms involves neuronal overexcitation induced by prolonged current flow, ultimately leading to excitotoxicity. Evidence for such mechanisms has been presented primarily for pulsed stimulation protocols [14]. We consider this explanation unlikely in the present study, as we employed cathodal iDCS, which under many conditions suppresses neuronal activity (e.g., [29]).

A second mechanism that could explain tissue damage involves Joule heating associated with current flow through tissue [30]. This is unlikely the primary cause of tissue damage for most of the smaller doses used in the present study because Zhang et al. [17] found that suprathreshold currents directly delivered to the intact dura for 15 minutes did not increase the brain surface temperature.

We therefore conclude that the neural injuries observed here are most likely caused by mechanisms related to the generation of toxic electrochemical reaction products at the electrode–tissue interface. Cathodal iDCS, used here, results in reductive processes involving electron transfer from the electrode to the tissue, leading to hydrogen gas formation, increased hydroxyl ion concentration and pH elevation, oxide formation, and hydrogen adsorption on platinum electrodes. For anodal iDCS, which was not used here, oxidative reactions and thus different electrochemical species prevail, possibly causing different neural injuries and other safety boundaries (see [18, 19]).

To derive a safety limit for iDCS, we followed the approach introduced by Liebetanz and colleagues [13] for tDCS. We quantified total lesion volume across all experiments and expressed stimulation dose as charge density, a parameter known to be relevant for describing neural injury caused by pulsatile stimulation [14]. Because Liebetanz et al. [13] reported a strong linear relationship between charge density and lesion size (*R*^2^ = 0.94), we likewise applied a linear regression model. This analysis yielded a somewhat lower, but still substantial, coefficient of determination (*R*^2^ = 0.68) and an estimated iDCS safety boundary of 2.3 mC/mm^2^. In the absence of a mechanistic basis favoring a specific dose–lesion relationship [31], we tested alternative mathematical models and found that power-law relationships with exponents greater than 1 modestly increased the *R*^2^ value (up to 0.77) and yielded less conservative safety values of up to 8.7 mC/mm^2^.

For the stimulation parameters used in this study, our approach yielded a charge density threshold of minimally 2.3 mC/mm^2^. This implies that it may be possible to safely apply a current of approximately 16 µA for 10 minutes to the cortex using electrodes with a surface area of approximately 4 mm^2^. This current generates an electric field of approximately 20 mV/mm at the brain surface, which decreases to about 10% of its peak value within a distance of a few millimeters. Therefore, iDCS could enable experimental paradigms and applications in animals and humans using stronger and more focal electric fields than with tDCS. In the latter case, currents of up to 2 mA (and occasionally up to 4 mA) are typically used, producing electric fields below 1 mV/mm in the brain [31]. Such field strengths are often considered too weak to produce robust effects, and have been considered possible explanation for the limited insights obtained from tDCS and the growing skepticism regarding its utility for studying and modulating human brain function [32].

Because the present estimates are expressed as total charge, higher currents and field strengths should be feasible for shorter stimulation durations, whereas longer durations would require lower current and field limits. For prolonged stimulation durations, the maximum safe current may in fact be higher than predicted by a simple cumulative charge model, because electrochemical byproducts at the electrode-tissue interface are likely to accumulate relatively slowly and may be partially cleared by diffusion and by interstitial fluid homeostasis mechanisms within the brain [16]. This possibility may explain why Hurlbert et al. [18] observed no histological damage in the spinal cord following the application of 1.5 µA DC over periods ranging from several hours to multiple days, even though the cumulative charge would theoretically exceed the threshold identified here after approximately 106 minutes. Because the kinetics of clearance mechanisms remain poorly characterized, it is currently unclear whether the safety limits for stimulation periods shorter than 10 minutes are smaller than those predicted by the present results. Likewise, it remains an open question whether the detrimental effects of electrochemical accumulation could be mitigated by intermittent stimulation with short stimulation pauses (“charge-balanced”) iDCS, in which cathodal and anodal stimulation phases alternate over periods of seconds. Lastly, alternative approaches for delivering ionic DC could be developed to avoid direct charge transfer through metal electrodes [6].

A possible application of iDCS could be neurocognitive research, where the use of the present stimulation parameters would enable performance of causal studies in animals or humans [33] to investigate the functional contribution of spatially confined cortical regions during perceptual or cognitive tasks. If the 10 minutes of continuous iDCS are divided into multiple shorter stimulation epochs lasting several seconds each, subjects could perform hundreds of behavioral trials while iDCS is applied.

An additional application of iDCS could exploit its suppressive effects in closed-loop systems for epilepsy control [34], where cathodal stimulation may provide an alternative means to terminate pathological brain activity locally, complementing existing approaches such as pulsed stimulation that aim to desynchronize neural activity.

In conclusion, the present study does not fully eliminate electrochemical concerns associated with DC delivery through metal electrodes. Nevertheless, it provides a quantitative starting point for further studies addressing electrochemical byproducts, chronic tissue responses, repeated stimulation, and functional consequences. In this sense, they may motivate renewed investigation of invasive DC stimulation, a largely underexplored field with physiological effects that differ from those of pulsatile stimulation.

## Acknowlegdements

We thank Douglas C. Fredericks for help during the experiments and the Comparative Pathology Laboratory in the Department of Pathology at the University of Iowa. This work was supported by grants of the National Institutes of Health (NIH/NIDCD R01 DC004290) and the Deutsche Forschungsgemeinschaft (DFG, project number 457346369).

## Declaration of competing interest

The authors declare that they have no competing financial interests or personal relationships that could be perceived to have influenced the work reported in this paper.

## Authorship Statement

Michael Brosch contributed to conceptualization, data curation, formal analysis, software, visualization, writing of the original draft, and review and editing of the manuscript. Hiroyuki Oya contributed to conceptualization, data curation, formal analysis, funding acquisition, investigation, methodology, project administration, resources, supervision, validation, and review and editing of the manuscript. Katherine N. Gibson-Corley contributed to conceptualization, data curation, formal analysis, investigation, methodology, resources, validation, visualization, and review and editing of the manuscript. Oliver Flouty contributed to conceptualization, data curation, formal analysis, investigation, methodology, resources, validation, and review and editing of the manuscript. Matthew A. Howard III contributed to conceptualization, funding acquisition, supervision, and review and editing of the manuscript. Kirill V. Nourski contributed to data curation, project administration, validation, and review and editing of the manuscript.

## Notes

### Competing Interest Statement

The authors have declared no competing interest.

## References

[1] Sutton AE, Krogmann RJ, Al Khalili Y. Cochlear implants. [Updated 2025 Jan 22]. In: StatPearls [Internet]. Treasure Island (FL): StatPearls Publishing; 2026 Jan-. Available from: https://www.ncbi.nlm.nih.gov/books/NBK544280/

[2] Benabid AL, Chabardes S, Mitrofanis J, Pollak P. Deep brain stimulation of the subthalamic nucleus for the treatment of Parkinson’s disease. Lancet Neurol. 2009;8:67–81. doi:10.1016/S1474-4422(08)70291-6.

[3] Stagg CJ, Antal A, Nitsche MA. Physiology of transcranial direct current stimulation. J ECT. 2018;34:144–52. doi:10.1097/YCT.0000000000000510.

[4] Jackson MP, Truong D, Brownlow ML, Wagner JA, McKinley RA, Bikson M, et al. Safety parameter considerations of anodal transcranial direct current stimulation in rats. Brain Behav Immun. 2017;64:152–61. doi:10.1016/j.bbi.2017.04.008.

[5] Merrill DR, Bikson M, Jefferys JG. Electrical stimulation of excitable tissue: design of efficacious and safe protocols. J Neurosci Methods. 2005;141:171–98. doi:10.1016/j.jneumeth.2004.10.020.

[6] Aplin FP, Fridman GY. Implantable direct current neural modulation: theory, feasibility, and efficacy. Front Neurosci. 2019;13:379. doi:10.3389/fnins.2019.00379.

[7] Zhao DX, Green AL, Purcell EK. Tissue response to deep brain stimulation electrodes: a review of animal and neurohistopathological studies. J Neural Eng. 2025. doi:10.1088/1741-2552/adf669.

[8] Kuncel AM, Grill WM. Selection of stimulus parameters for deep brain stimulation. Clin Neurophysiol. 2004;115:2431–41. doi:10.1016/j.clinph.2004.05.031.

[9] Cogan SF, Ludwig KA, Welle CG, Takmakov P. Tissue damage thresholds during therapeutic electrical stimulation. J Neural Eng. 2016;13:021001. doi:10.1088/1741-2560/13/2/021001.

[10] Shannon RV. A model of safe levels for electrical stimulation. IEEE Trans Biomed Eng. 1992;39:424–6. doi:10.1109/10.126616.

[11] Ackermann DM Jr, Bhadra N, Foldes EL, Kilgore KL. Separated interface nerve electrode prevents direct current induced nerve damage. J Neurosci Methods. 2011;201:173–6. doi:10.1016/j.jneumeth.2011.01.016.

[12] Fridman GY, Della Santina CC. Safe direct current stimulation to expand capabilities of neural prostheses. IEEE Trans Neural Syst Rehabil Eng. 2013;21:319–28. doi:10.1109/TNSRE.2013.2245423.

[13] Liebetanz D, Koch R, Mayenfels S, König F, Paulus W, Nitsche MA. Safety limits of cathodal transcranial direct current stimulation in rats. Clin Neurophysiol. 2009;120:1161–7. doi:10.1016/j.clinph.2009.01.022.

[14] McCreery DB, Agnew WF, Yuen TGH, Bullara L. Charge density and charge per phase as cofactors in neural injury induced by electrical stimulation. IEEE Trans Biomed Eng. 1990;37:996–1001. doi:10.1109/10.102812.

[15] Chhatbar PY, George MS, Kautz SA, Feng W. Quantitative reassessment of safety limits of tDCS for two animal studies. Brain Stimul. 2017;10:1011–2. doi:10.1016/j.brs.2017.07.008.

[16] Hamm LL, Nakhoul N, Hering-Smith KS. Acid-base homeostasis. Clin J Am Soc Nephrol. 2015;10:2232–42. doi:10.2215/CJN.07400715.

[17] Zhang K, Guo L, Zhang J, An G, Zhou Y, Lin J, et al. A safety study of 500 μA cathodal transcranial direct current stimulation in rat. BMC Neurosci. 2019;20:40. doi:10.1186/s12868-019-0523-7.

[18] Hurlbert RJ, Tator CH, Theriault E. Dose-response study of the pathological effects of chronically applied direct current stimulation on the normal rat spinal cord. J Neurosurg. 1993;79:905–16. doi:10.3171/jns.1993.79.6.0905.

[19] Horsley V, Clarke RH. The structure and functions of the cerebellum examined by a new method. Brain. 1908;31:45–124. doi:10.1093/brain/31.1.45.

[20] Hughes GB, Bottomy MB, Dickins JR, Jackson CG, Sismanis A, Glasscock ME 3rd. A comparative study of neuropathologic changes following pulsed and direct current stimulation of the mouse sciatic nerve. Am J Otolaryngol. 1980;1:378–84. doi:10.1016/s0196-0709(80)80018-4.

[21] Hughes GB, Bottomy MB, Jackson CG, Glasscock ME 3rd, Sismanis A. Myelin and axon degeneration following direct current peripheral nerve stimulation: a prospective controlled experimental study. Otolaryngol Head Neck Surg. 1981;89:767–75. doi:10.1177/019459988108900515.

[22] Bray IE, Clarke SE, Casey K, Nuyujukian P, Brain Interfacing Laboratory. Neuroelectrophysiology-compatible electrolytic lesioning. Elife. 2024;12. doi:10.7554/eLife.84385.2.

[23] Rathi S, Krzemiński J, Vighneshvel T, Pepłowski A, Baraniecki D, Tiwari N, Karpova A, Janczak D, Jakubowska M and Brosch M (2026) Biological performance evaluation of graphene nanoplatelets for intracranial direct current stimulation. Front. Neurosci. 20:1664084. doi: 10.3389/fnins.2026.1664084.

[24] Nourski KV, Howard MA 3rd. Invasive recordings in the human auditory cortex. Handb Clin Neurol. 2015;129:225–44. doi:10.1016/B978-0-444-62630-1.00013-5.

[25] Gibson-Corley KN, Oya H, Flouty O, Fredericks DC, Jeffery ND, Gillies GT, et al. Ovine tests of a novel spinal cord neuromodulator and dentate ligament fixation method. J Investig Surg. 2012;25:366–74. doi:10.3109/08941939.2012.677967.

[26] Flouty OE, Oya H, Kawasaki H, Reddy CG, Fredericks DC, Gibson-Corley KN, et al. Intracranial somatosensory responses with direct spinal cord stimulation in anesthetized sheep. PLoS One. 2013;8. doi:10.1371/journal.pone.0056266.

[27] Knight RD. Physics for Scientists and Engineers: A Strategic Approach with Modern Physics. 5th ed. Hoboken: Pearson; 2022.

[28] Khatoun A, Asamoah B, Mc Laughlin M. Investigating the feasibility of epicranial cortical stimulation using concentric-ring electrodes: a novel minimally invasive neuromodulation method. Front Neurosci. 2019;13:773. doi:10.3389/fnins.2019.00773.

[29] Bikson M, Inoue M, Akiyama H, Deans JK, Fox JE, Miyakawa H, et al. Effects of uniform extracellular DC electric fields on excitability in rat hippocampal slices in vitro. J Physiol. 2004;557:175–90. doi:10.1113/jphysiol.2003.055772.

[30] Bikson M, Datta A, Elwassif M. Establishing safety limits for transcranial direct current stimulation. Clin Neurophysiol. 2009;120:1033–4. doi:10.1016/j.clinph.2009.03.018.

[31] Soleimani G, Alekseichuk I, Aurup C, Bergmann TO, Bestmann S, Beynel L, et al. Dose-response relationships in transcranial brain stimulation: physics, physiology and mechanism. Brain Stimul. 2026;19:103067. doi:10.1016/j.brs.2026.103067.

[32] Filmer HL, Mattingley JB, Dux PE. Modulating brain activity and behaviour with tDCS: rumours of its death have been greatly exaggerated. Cortex. 2020;123:141–51. doi:10.1016/j.cortex.2019.10.006.

[33] Nourski KV, Steinschneider M, Rhone AE, Howard MA 3rd. Stimulus-driven and behavior-driving activity along the cortical auditory hierarchy. NeuroImage. 2026;328:121801. doi:10.1016/j.neuroimage.2026.121801.

[34] Nair DR, Laxer KD, Weber PB, Murro AM, Park YD, Barkley GL, et al.,,,,, RNS System LTT Study. Nine-year prospective efficacy and safety of brain-responsive neurostimulation for focal epilepsy. Neurology. 2020;95–56. doi:10.1212/WNL.0000000000010154.

